# A miniaturised semi-dynamic *in-vitro* model of human digestion

**DOI:** 10.1101/2024.04.17.589902

**Authors:** Victor Calero, Patrícia M. Rodrigues, Tiago Dias, Alar Ainla, Adriana Vilaça, Lorenzo Pastrana, Miguel Xavier, Catarina Gonçalves

**Affiliations:** International Iberian Nanotechnology Laboratory, Avenida Mestre José Veiga, s/n, 4715-330, Braga, Portugal; Nova School of Science and Technology, Nova University of Lisbon, Lisbon, Portugal

## Abstract

Reliable *in-vitro* digestion models that are able to successfully replicate the conditions found in the human gastrointestinal tract (GIT) are key to assess the fate and efficiency of new formulations aimed for oral consumption. However, current *in-vitro* models either lack the capability to replicate crucial dynamics of digestion or require large volumes of sample/reagents, which can be scarce when working with nanomaterials under development. Here, we propose a miniaturised digestion system, a digestion-chip, based on incubation chambers integrated on a polymethylmethacrylate (PMMA) device. The digestion-chip incorporates key dynamic features of human digestion, such as gradual acidification and gradual addition of enzymes and simulated fluids in the gastric phase, and controlled gastric emptying, while maintaining low complexity and using small volumes of sample and reagents. In addition, the new approach integrates real-time automated closed-loop control of two key parameters, pH and temperature, during the two main phases of digestion (gastric and intestinal) with an accuracy down to ±0.1°C and ±0.2 pH points. The experimental results demonstrate that the digestion-chip successfully replicates the gold standard static digestion INFOGEST protocol and that the semi-dynamic digestion kinetics can be reliably fitted to a first kinetic order model. These devices can be easily adapted to dynamic features in an automated, sensorised, and inexpensive platform and will enable reliable, low-cost and efficient assessment of the bioaccessibility of new and expensive drugs, bioactive ingredients or nano-engineered materials aimed for oral consumption, thereby avoiding unnecessary animal testing.

## Introduction

Human digestion is a complex multistage process, where ingested compounds undergo a wide variety of physical and biochemical steps before reaching the intestinal epithelium for absorption^1^. Ingested compounds are exposed to a harsh environment, which might cause their degradation/disintegration throughout the gastrointestinal tract (GIT) and, therefore affect their bioaccessibility, i.e., the fraction that reaches the small intestine and is available for absorption^2^. In addition, these compounds must undergo intestinal absorption to reach systemic circulation and become bioavailable^3^.

As an alternative to *in-vivo* models, *in-vitro* digestion models are attracting growing interest as a mean to avoid the challenges associated to animal testing, aiming to increase reliability and reduce failures in human clinical trials^1,4,5^. These models are usually categorised as static or dynamic^6–8^.

In static digestion models, the different steps of human digestion are mimicked inside closed reservoirs keeping the digestive conditions constant over the incubation period. During each phase, the sample is mixed with simulated digestive fluids and incubated for a specific time at constant temperature, enzyme activity and pH. In 2014, the INFOGEST network proposed a standardised protocol to minimise variability of the experimental conditions used to emulate human digestion in *in-vitro* models described elsewhere^9–11^, allowing read-across of experimental results between different research teams^10,12^. The INFOGEST protocol defines a sequential oral, gastric and intestinal digestion process where pH, incubation times, type and activity of enzymes, bile concentration, sample dilution and simulated fluid composition were defined based on available physiological data^13^. Some studies have validated the convenience of static digestion models due to their simplicity and reduced costs, and indicated that they can provide a good end-point accuracy^1,10,14^. In many cases, they seem too simplistic to deliver reliable results, mainly when kinetics are pertinent to the study. This has been extensively described in several reviews^4,7,15,16^.

In contrast, dynamic models incorporate relevant features to replicate the complexity of the digestion process.^16,17^ These include continuous flow, controlled addition of enzymes and simulated fluids, monitoring and automatic adjustment of pH, peristalsis, and gastric emptying. Thus, *in-vitro* dynamic models provide a better approximation to the conditions found *in-vivo*, enable time-resolved analysis, and provide results that can be directly correlated with *in-vivo* studies^18^. There are some examples found in literature, such as the TNO gastrointestinal model (TIM)^19^ and the Simulator of the Human Intestinal Microbial Ecosystem (SHIME) model^20^, which combine physiological processes within the stomach and the intestine providing important information regarding the digestibility of food matrices. These models are multi-compartmental, meaning that they contain different compartments for the different parts of the GIT and include real-time monitoring of different variables, such as pH and temperature. Another example is the Gastrointestinal Simulator developed at the Institute of Food Science Research (CIAL, Spain)^21^, which has recently been used to understand the potential effects of nanoparticles on the simulated digestive tract.

Nevertheless, these sophisticated models are complex, time-consuming and require large amounts of reagents and enzymes^1^, constituting a major drawback when applied to test expensive nanomaterials or drugs. As a compromise, Mulet-Cabero *et al*. ^17^from the INFOGEST network proposed a semi-dynamic protocol that, while following harmonised protocols, fills the gap between the reliable but complex dynamic models and the over simplistic but affordable static models. It is named semi-dynamic as it includes dynamics only in the gastric phase keeping the intestinal phase totally static (2h of incubation using constant conditions: pH and enzyme activities). In the gastric phase, the semi-dynamic protocol simulates the gradual addition of gastric secretions, gradual acidification and performs several gastric emptying, enabling the assessment of the role of these dynamics in the digestion of ingested products. The protocol does not recommend specific apparatus, as an example an auto-titrator including a pH probe and a dosing unit or alternatively a syringe pump with a pH probe, are mentioned. The temperature must be controlled using a vessel with a thermostat jacket or other approach.

Here, we present a novel miniaturised system for *in-vitro* digestion studies. This versatile platform can be easily adapted to a static or a semi-dynamic protocol, incorporating the key dynamics mentioned above, in the gastric phase, mimicking the transient nature of gastric secretions and emptying. In this regard, our approach focuses on the upper parts of the digestive tract (mouth, stomach and small intestine), alike the DIDGI® system developed at the French National Institute for Agricultural research (INRA)^22,23^.

Nevertheless, our miniaturised digestion device brings several benefits over the current *in-vitro* digestion models. There is a significant reduction of the footprint of the experimental setup when compared to the current dynamic models and minimal user input is required, given its automated operation. The simplicity of the device allows for easy sampling during the digestion simulation and continuous monitoring ensures that pH and temperature are automatically controlled during the experiment. Finally, the small dimensions of the incubation chambers translate into low consumption of sample/reagent volumes.

Finally, it is important to note that the choice of an appropriate *in-vitro* digestion model depends on the research question being addressed. In some cases, static models may be sufficient to provide accurate end-point digestion assessments. More complex and dynamic models may be necessary to accurately replicate digestion and evaluate the physicochemical changes certain foods undergo along the digestive tract. While our miniaturised system may not be suitable for *in-vitro* digestion of certain food matrices, it will become particularly attractive to evaluate the bioaccessibility of nano-encapsulated expensive drugs or bioactive ingredients, which are often produced in limited quantities during the development stage^24^.

## Results and discussion

### Temperature and pH control

Before the experiments, the two pairs of pH electrodes (sensing & reference) were calibrated using a four-point calibration by sequentially immersing the probes in standard pH reference solutions, with pH = 1.7, 4.0, 7.0 and 9.2.

In Electronic Supplementary Information (ESI), figure S1a shows the response of the potential difference to the calibration solutions, showing a fast response and stabilisation (under 60 seconds) of the sensors. The calibration plots are displayed in Figure S1b, showing a good linear response. Slope values typically ranged between –65 mV/pH and –70 mV/pH. The performance of these probes was then further validated by comparing the pH measurements to a commercial pH-meter (Mettler Toledo®) in SGF solutions with different pH values ranging from 2.0 to 7.0. Maximum discrepancies were found with around ± 0.2 pH points for the more acidic pH values (2.0 to 3.0) and were minimal in the range of pH 3.0 to 7.0 (under ± 0.1 pH points). Furthermore, the probes long-term stability was validated in SGF (pH 3.0), showing a ± 10 mV variation over a time period exceeding 10 hours. The stability of the electrode pair and of each electrode independently against a commercial probe can be found in ESI (Figure S2).

Figure 1 shows an example of the temperature and pH monitoring and control in a (a) static digestion using only one digestion chamber – and thus with a single pH probe and a temperature sensor – and (b) semi-dynamic digestion using both incubation chambers and two pairs of sensors. Figure 1a shows the temperature (in red) in the chambers, which is set at 37 °C and maintained constant throughout the entire experiment. The spikes around 150 minutes are due to the addition of the intestinal components (SIF + Pancreatin + Bile salts) to the digestion chamber. The pH (in blue) is first adjusted from a value around 6.5 to 3.0 for the gastric phase of digestion. Then, the pH is increased back to 7.0 for the intestinal phase of digestion. In this example, the pH reached a value higher than the set value of 7.0 due to an excessive addition of 1M NaOH. This was mitigated by the subsequent addition of 1M HCl, showing the versatility and reliability of the platform. Figure 1b.1 shows the pH data from a semi-dynamic digestion. The pH in the gastric chamber is gradually reduced over the course of the digestion, whereas the pH in the intestinal chamber is maintained at a pH value around 7.5. Figure 1b.2 shows the temperature readings from the same experiment.

**Figure 1.**
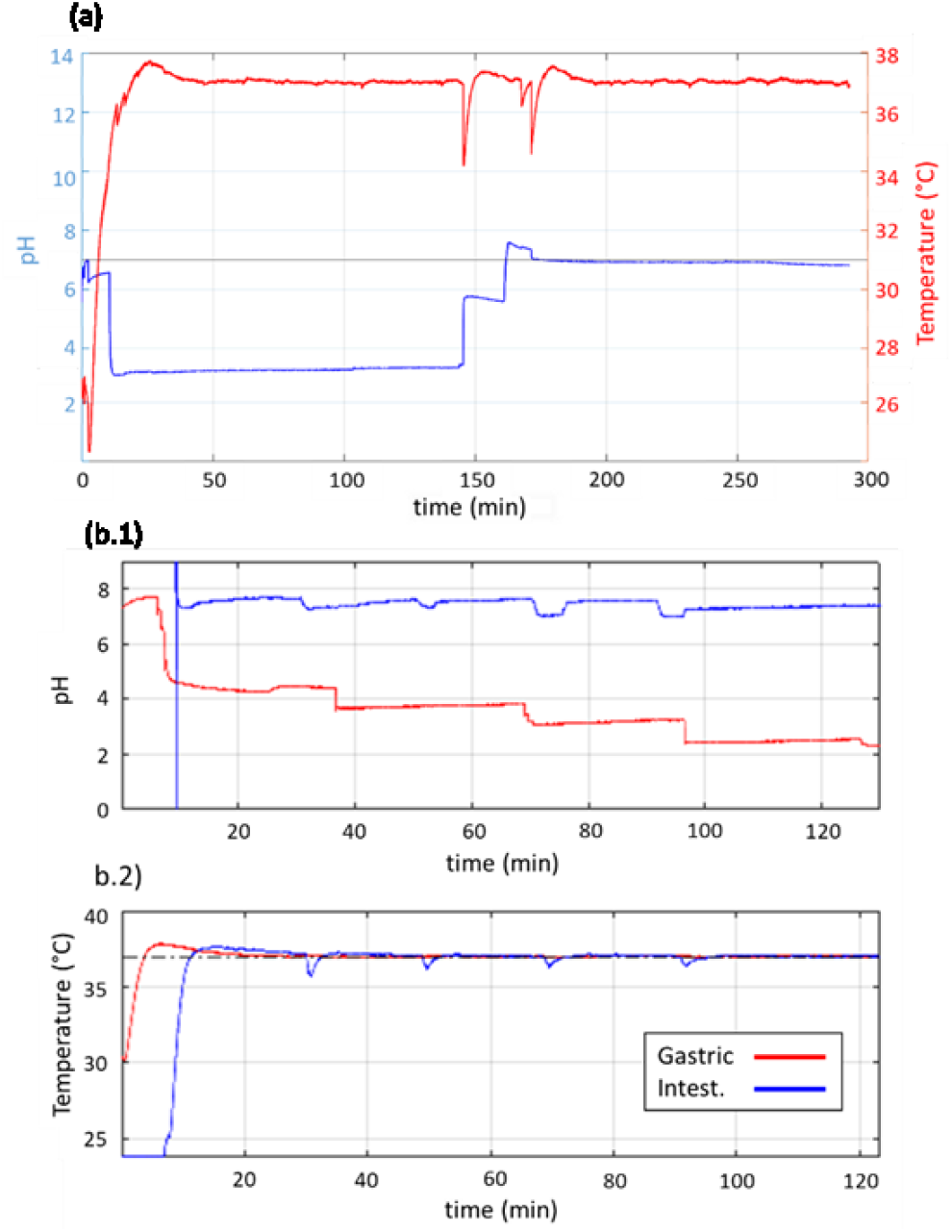
(a) Example of real-time temperature and pH control during a static digestion experiment in a single incubation chamber. The temperature of the digestive fluids is maintained constant at 37 ºC while the pH is first adjusted to 3.0 in the gastric phase and then increased to 7.0 in the intestinal phase. (b.1) Example pH data from a semi-dynamic digestion showing gradual acidification of the gastric phase (in red) while the intestinal phase (in blue) is kept at a constant pH around 7-7.5. (b.2) Temperature data corresponding to the gastric and intestinal incubation chambers. The spikes observed in the intestinal phase correspond to each gastric emptying where the SIF is introduced at room temperature and mixed with the gastric phase (at 37 ºC).

### Static *in-vitro* digestion

To validate static *in-vitro* digestions using the digestion-chip, we replicated the INFOGEST protocol in our device and compared the results with the standard bench-top approach. Figure 2 shows the extension of digestion of the two test molecules in terms of the normalised fluorescence intensity measured and as a function of time. The fluorescence intensity measured at each wavelength is proportional to the extension of hydrolysis of the respective substrate. Figure 2a shows the digestion of the protease substrate, which expectedly increased rapidly at the start of each digestion phase when the concentration of substrate [S] was high and the concentration of product [P] was low, to then reach a plateau showing a completed hydrolysis by pepsin, the protease of the gastric phase. This could be anticipated given that pH and enzyme concentration [E] were constant throughout both digestion phases. This translates into a constant enzyme activity, which in the case of pepsin acting in the gastric phase is optimal within the pH range 2.0-3.0. In the case of the pancreatic enzymes acting in the intestinal phase, it is optimal within pH 6.0-8.0. The digestion on-chip replicates the time-dependence obtained during the standard bench-top protocol.

**Figure 2.**
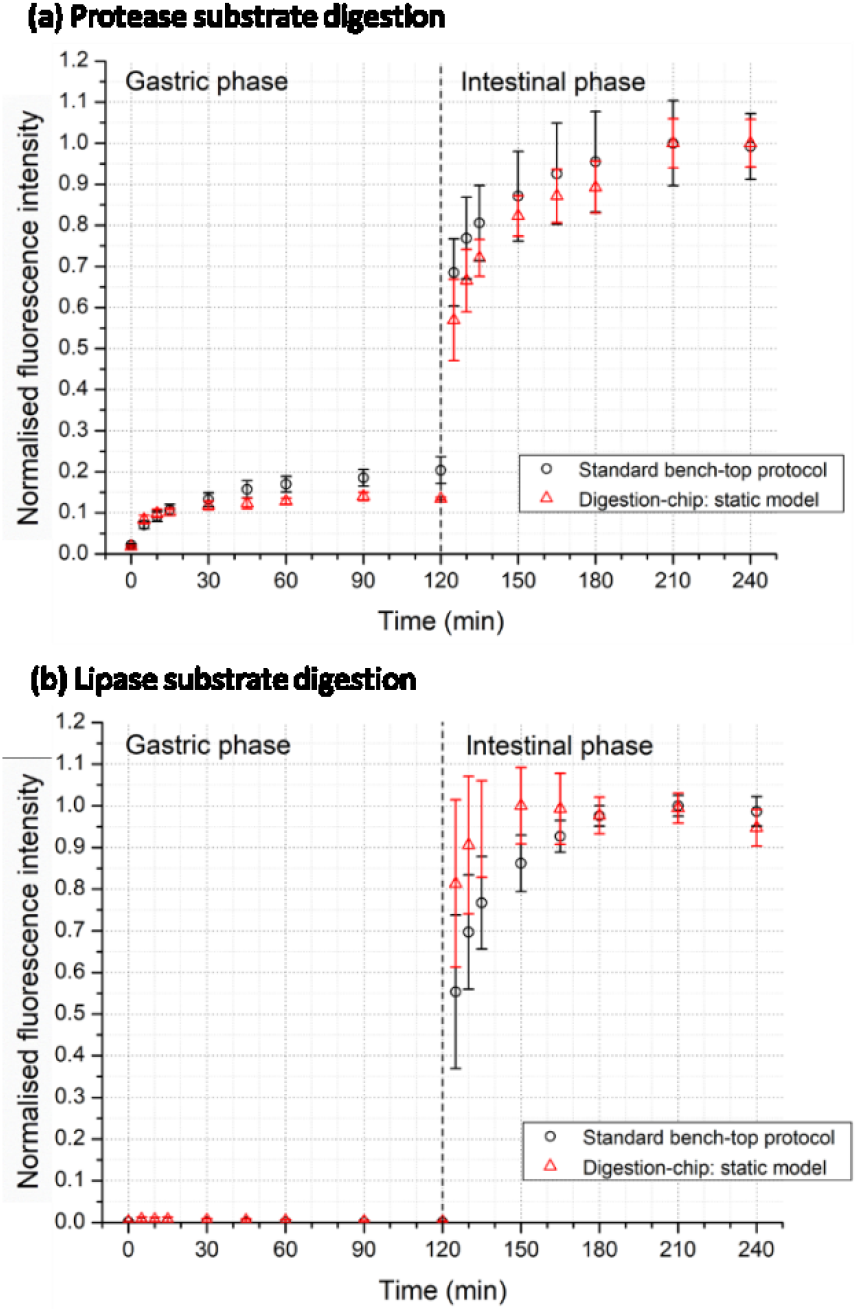
Comparison of the static in-vitro digestion in the digestion-chip (N=3) and the standard bench-top protocol (N=4), both following the INFOGEST protocol. The data is presented as normalised fluorescence intensity, which is proportional to the concentration of hydrolysed protease and lipase substrates. (a) Protease substrate digestion (λex = 589 nm; λem = 617 nm). (b) Lipase substrate digestion (λex = 482 nm; λem = 515 nm).

Figure 2b shows the digestion of the lipase substrate in equivalent experiments. This substrate is not digested during the gastric phase, due to the absence of lipase enzymes, as shown by the nearly null values measured during the whole duration of the gastric phase. However, when the lipase substrate enters the intestinal phase of digestion, there is a rapid increase due to the high amount of substrate available and the action of lipases present in pancreatin. Here, the digestion was slightly faster for the digestion-chip and then reaches a plateau following the trends observed in the standard bench-top protocol.

The results summarised in Figure 2 demonstrate the capabilities of this experimental setup to replicate digestive conditions, with an automated control of the experimental conditions while using low volumes of sample and reagents. Our device circumvents the need to do preliminary tests to assess the volumes of acid and base needed to adjust pH and provides its constant monitoring, together with temperature, thus avoiding adjustment errors. In addition, these results enabled to discard possible enzyme degradation due to mechanical strain or localised excessive heating near the heating elements.

### Semi-dynamic in-vitro digestion

Nevertheless, as previously discussed, static digestion protocols fail to replicate the complexity of human digestion. An advantage of the digestion-chip is that it allows a very simple integration of key dynamic digestion features, while maintaining a small footprint and using low sample and reagent volumes. To validate the capacity of the device to integrate dynamic features in the gastric phase we did experiments including gradual acidification, gradual addition of enzymes and gastric emptying. The results from the on-chip semi-dynamic *in-vitro* digestions are summarised in Figure 3, where the experimental data is normalised to the highest measured fluorescence value and the concentration of digestion products is assumed to be proportional to the fluorescence intensity.

**Figure 3.**
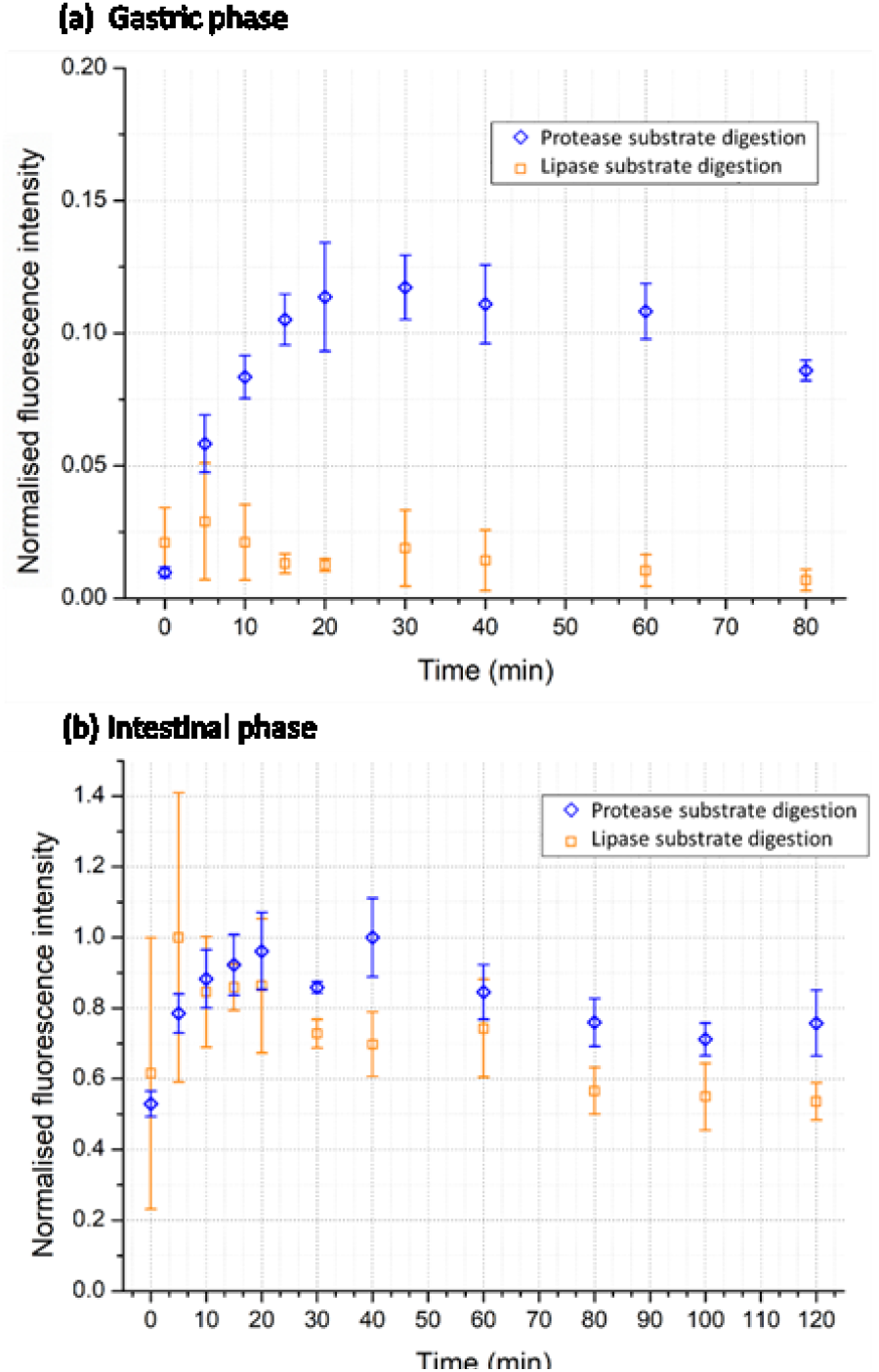
Normalised fluorescent intensity from the semi-dynamic in-vitro digestion of protease and lipase substrates (N=3). The extension of digestion is proportional to the fluorescence intensity measured from the hydrolysed protease and lipase substrates. (a) Gastric phase. (b) Intestinal phase.

Figure 3a shows the digestion of the protease and lipase substrates during the gastric phase. As expected, there is a more gradual increase in the protein digestion when compared to the static conditions (Figure 2a) given the linear dependence with time for the first 20 minutes of the semi-dynamic digestion. This is due to the gradual acidification of the gastric phase (see Figure 1b.1) and the gradual addition of pepsin. The product concentration then stabilises and starts to decrease reaching owing to a completed hydrolisation of the substrate from the gastric enzymes (pepsin) and the dilution from the incoming gastric fluids. Concurrently, lipid digestion was not significant since lipases are not included in the gastric phase. This is reflected by the very low fluorescence intensity shown in Figure 3a.

For intestinal digestion, 4 gastric emptying were performed at time-points 20, 40, 60 and 80 minutes. Additionally, an extra emptying was done at time-point 0 (without gastric incubation time) to analyse the kinetics of the intestinal enzymes in isolation. Figure 3b shows that there was a rapid increase in the extension of digestion of both substrates, similar to the static experiments. The digestion of the protease substrate occurs fast following the first emptying to then slowly decrease until it stabilises close to the end of the incubation. The lipase substrate digestion follows the same trend as the protease substrate, given that enzyme concentration and pH are kept constant throughout the experiment.

In order to further validate the results from the on-chip semi-dynamic digestions, we fitted the protease digestion data (which shows clearer and richer dynamics) to a first order kinetics reaction model:

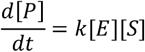

where *k* is the reaction rate constant. The model data is equally normalised to the highest digestion products concentration. We use a simple first order kinetics model, which is limited to a specific reaction. In this case, we have a complex substrate with different cleavage points. Thus, we can only define effective values for the rate constant *k*, since mechanistically it does not correspond to a specific hydrolysis reaction but to a sum of all different cleaving reactions.

Figure 4a shows the experimental results of the gastric phase together with the best model fitting (for *k* = 0.0043 *s*^−1^(*U*/μ*L*)^−1^). The model reproduces accurately the dependence observed in the gastric phase, with a sharp initial increase in product concentration followed by a slow decrease due to sample dilution once the completed hydrolysis by pepsin. A diagrammatic depiction of the evolution of [S], [E] and the total reaction volume is also included.

**Figure 4.**
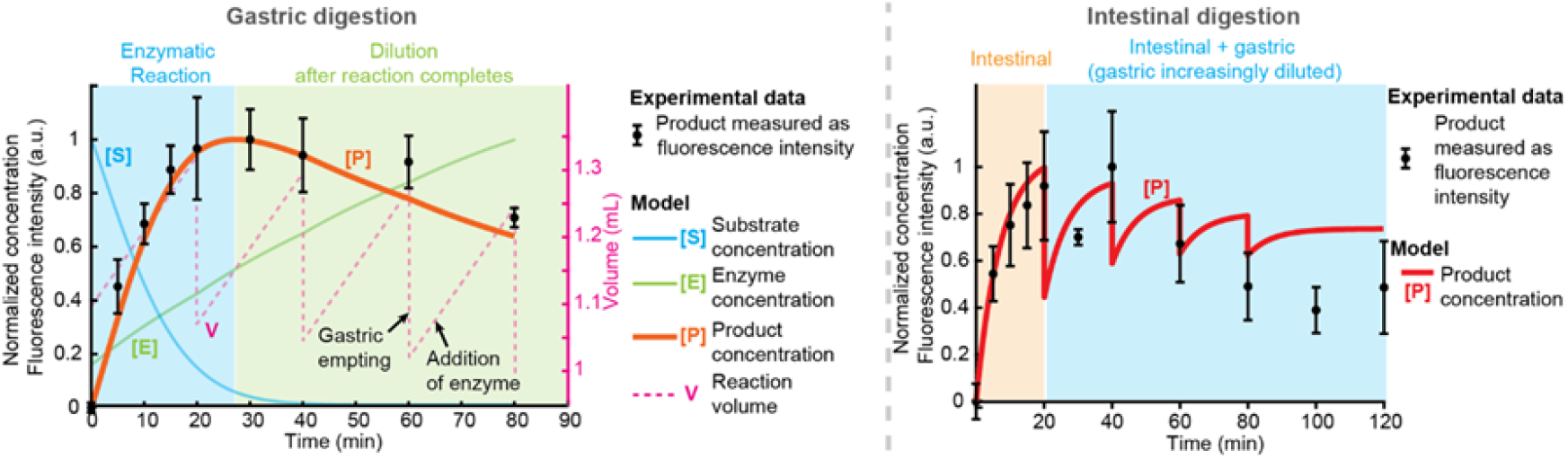
Results of the first-order kinetics model fitted to the experimental protease substrate digestion results in the digestion-chip. Best fitting of the model (red) for the digestion rate constant *k* to the experimental data from the semi-dynamic digestions (black). (a) Gastric phase, (b) intestinal phase.

The intestinal phase is much more complex (Figure 4b). Pancreatin is a mixture of enzymes, and intestinal digestion occurs on top of a partially digested substrate from the different gastric emptying, which are sequentially added to the same reaction chamber. Nevertheless, the model can predict well the first half of the intestinal phase (with *k* = 0.046 *s*^−1^(*U*/μ*L*)^−1^), showing a sharp increase at the start of the reaction after the first emptying. This is followed by a moderate decrease at t = 20 minutes, attributed to the dilution caused by the second gastric emptying. Another sharp increase follows, which justified by the continuation of the digestion.

The model predicts a sequential increase on the total digestion products concentration resulting from the cleavage of protease substrate by intestinal enzymes, which is restarted after each gastric emptying. To observe a better match with the experimental results, it would be necessary to increase the time resolution of the experiments. Finally, the model does not reproduce the decrease in product concentration to the same extent observed experimentally during the second half of the intestinal phase.

We hypothesise that this discrepancy could be due to enzyme degradation, possibly caused by auto-proteolytic activity, and enzyme product inhibition, which have been described in the literature^25–30^ but were not considered in our model. These would effectively reduce enzyme activity over time and therefore the total product content cannot account for the dilution caused by the increase in volume after each gastric emptying. In fact, the last gastric emptying occurs at t = 80 minutes, after which the intestinal product concentration remains constant until the end of the digestion, showing that there is no degradation of the product. In our devices all gastric emptying volumes are combined into the same intestinal phase and thus, the intestinal phase kinetics should be taken as an end-point measurement. Adding to our digestion-chip the capacity to incubate intestinal phase reactions independently is one of our current research goals.

## Conclusions

This work describes a novel *in-vitro* digestion platform, a digestion-chip, where digestion occurs inside miniaturised and automated incubation chambers with integrated sensors. The combination of an integrated peristaltic micro-pump and custom-made small-footprint syringe pumps, which are low-cost and programmable, allows sample and reagents to be automatically manipulated in the system. The continuous sensing of pH and temperature enables full automated control via feedback-loops driving the heating elements and the pumping of acid or base. The fabrication process is simple and affordable and the device design facilitates sampling during the digestion experiments.

The experimental results demonstrate that the digestion-chip can reliably perform static *in-vitro* digestions, replicating the gold-standard INFOGEST protocol while bringing several advantages, such as the automated adjustment of pH and temperature in each digestion phase. Additionally, the digestion-chip allows the integration of crucial dynamics of human digestion, providing more physiologically relevant outcomes through its use in semi-dynamic digestion simulations, and showing the capability to adapt to different protocols depending on the study requirements.

Furthermore, the reduced dimensions and low complexity of the setup, the small amounts of sample and reagents required and its user-friendly operation, make this approach particularly attractive for studying new drugs, bioactive compounds and nano-formulations. The production of such compounds is typically expensive during the early stages of development and requires extensive *in-vivo* testing to study their fate along the gastrointestinal tract^31^.

The choice of a correct in-vitro digestion model for each specific case-study is critical for an adequate assessment of the food digestibility and bioaccessibility and to guarantee the validity and applicability of the experimental outcomes. Our system allows for low-cost, facile and reliable assessment of bioaccessibility and digestion dynamics. We believe that this is a first prototype of a future family of versatile and miniaturised digestion simulation tools that can easily adapt diverse protocols, particularly useful for *in-vitro* digestion of new and expensive nanomaterials. Finally, coupling with gut-on-chip^32,33^ or multi-organ-chip models will enable replicating the complete gastrointestinal tract and analyse compound cytotoxicity, intestinal permeability and potentially first-pass metabolism, providing comprehensive studies of bioaccessibility and bioavailability in a single platform.

## Methods

### Device design and fabrication

A technical diagram of the digestion-chip can be found in Figure 5a, together with a description of the different elements integrated. The device comprises three circular compartments of 18 mm diameter that are two incubation chambers and one reservoir: 1) chamber to emulate the oral and gastric phase, 2) chamber to emulate the intestinal phase and 3) reservoir for simulated intestinal fluids, that will be added along the digestion process. As can be seen in figure 6, both gastric chamber and reservoir of simulated intestinal fluids are connected to the intestinal chamber by two peristaltic micro-pumps that pump simultaneously, at the same flow rate (1:1 ratio), “chyme” and the simulated intestinal fluids from the gastric chamber and the reservoir, respectively, into the intestinal chamber where intestinal phase is emulated. These devices were fabricated from PMMA sheets using CNC micro-milling (Flexicam Viper 606 with ArtCam software) and laser cutting (Widlaser LS1390 Plus with LaserWorkV6 software). The devices were assembled bonding several layers using double sided 3M tape (467MP; 3M, Saint Paul, MN, US).

**Figure 5.**
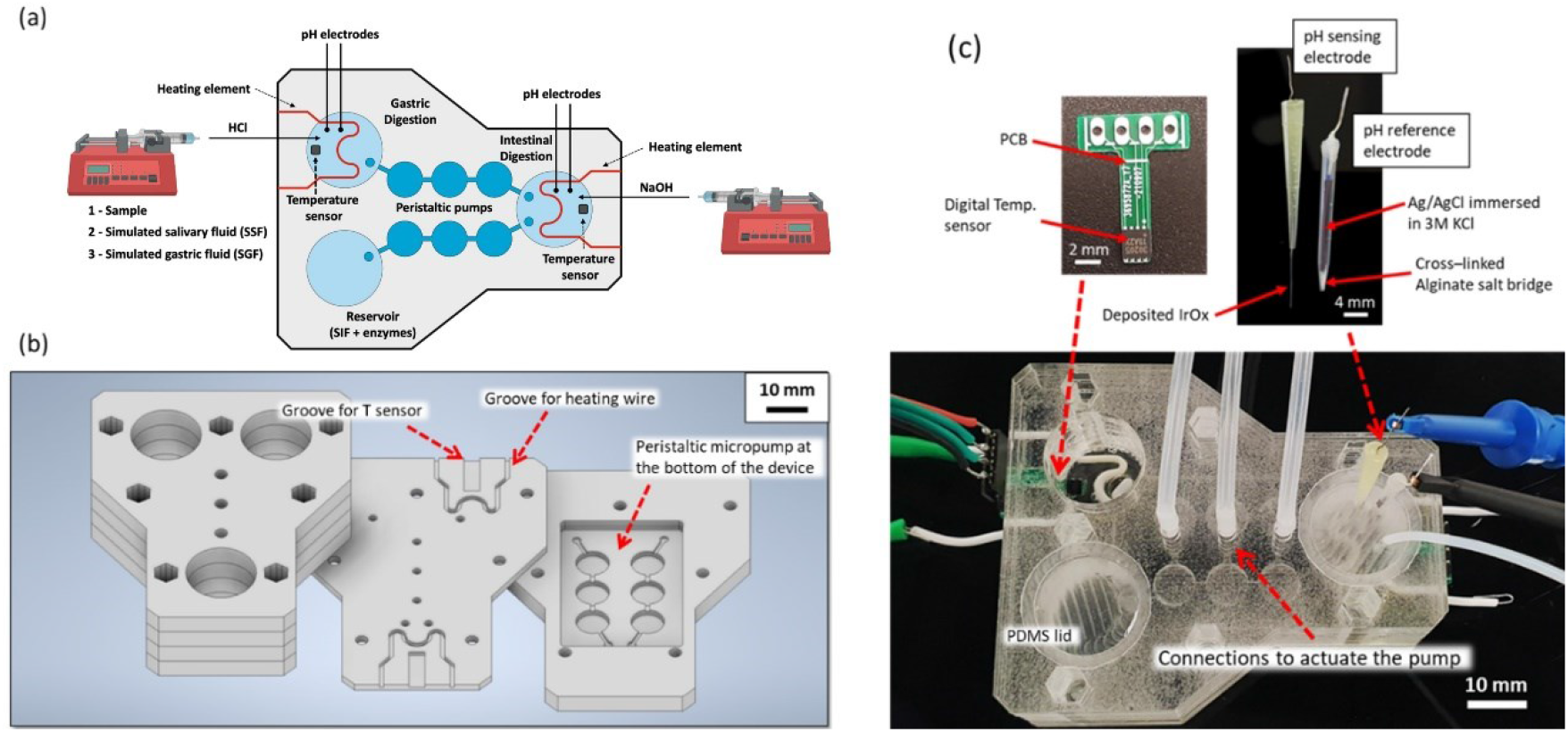
Technical description of the digestion-chip. (a) Diagram of the digestion chambers with the integrated components. (b) Description of the different layers that constitute the device. (c) Picture of the pH and temperature sensors and the digestion-chip.

**Figure 6.**
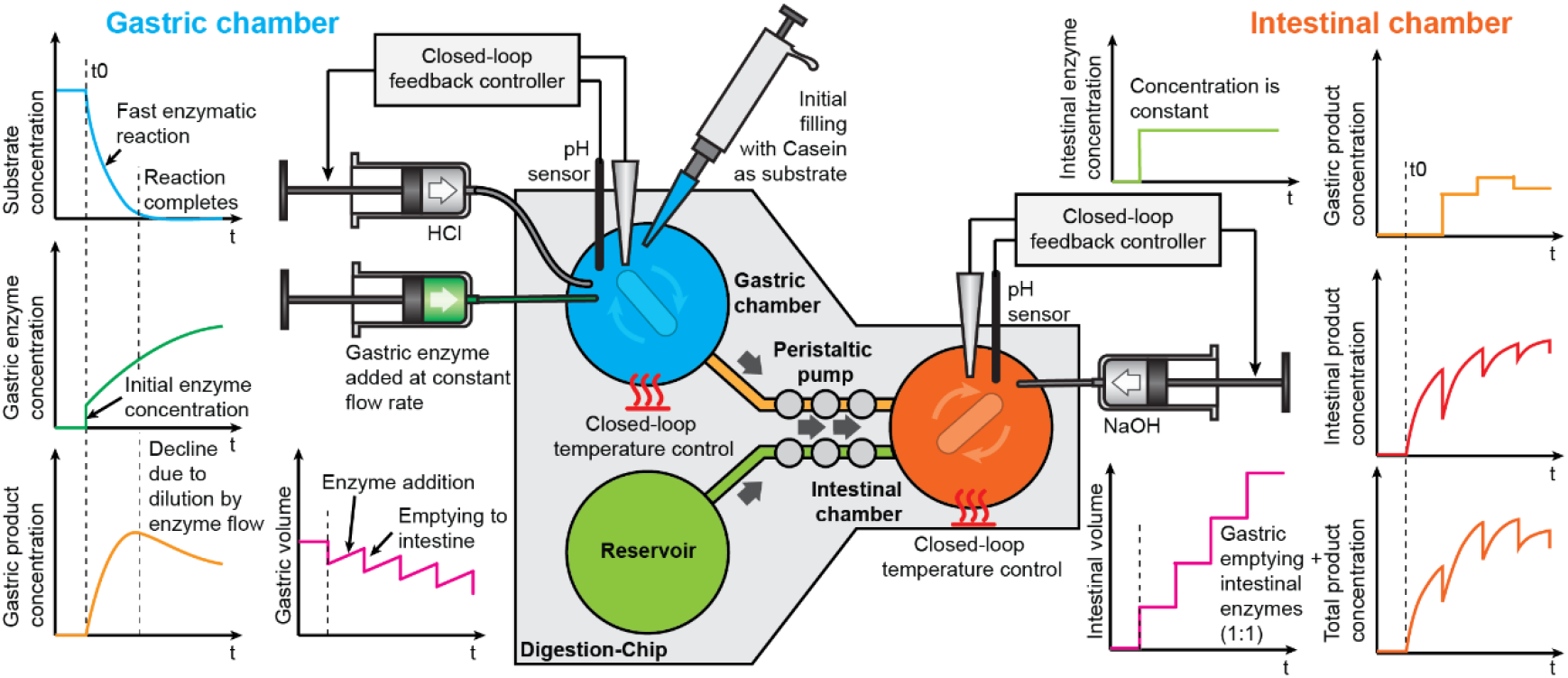
Conceptual diagram of the semi-dynamic digestion process in the digestion-chip. The graphs describe the evolution of sample volumes and of the concentration of substrate, reaction products and enzymes.

As shown in figure 5b, the device contains 6 layers: i) 4 layers (5 mm thickness) that define the height of the digestion chambers and reservoir, ii) 1 layer (3 mm thickness) that is the bottom layer of chambers and reservoir with grooves to accommodate temperature sensors and heating elements and iii) 1 bottom layer (5 mm thickness) to incorporate the peristaltic micro-pumps that allow gastric emptying. The bottom PMMA layer contains a rectangular cavity to place a PDMS piece with 2 parallel channels along with 3 chambers each bonded to a PDMS membrane. Here, the 6 chambers can be pressurised using a Fluigents MFCS™-EZ Pressure controller that bends the membrane, displacing the fluid between chambers. The bottom layer is not bonded but rather screwed to the PMMA layers. This fully seals the PDMS piece between the middle and bottom layers of PMMA and avoids leakage through the holes connecting the digestion chambers, reservoir, and the micro-pump channels. An image of the assembled device with all components integrated can be seen in Figure 5c. The pump is capable of driving up to 170 μL./min-1 and allows a precise control of the gastric emptying volume with approximately 50 μL per pumping cycle. A depiction of the pumping cycles is provided in Figure S3.

To ensure a constant mixing of the sample and the simulated digestive fluids, 2&5 mm magnetic stirring bars were placed at the bottom of the digestion chambers. The devices were then placed on top of a custom-made magnetic stirrer fabricated out of two pulse width modulation (PWM) controllable motors (recycled from computer cooling fans) placed below the digestion chambers. This provides efficient mixing without causing enzyme degradation by mechanical stress. Mixing is critical for these experiments, not only for enzyme-substrate reactions, but also to prevent localised fluid overheating near the heating elements (which could cause irreversible enzyme denaturation) and ensure fast pH equilibration. A picture of the custom magnetic stirrer and stirring bars can be found in the Electronic Supplementary Information (ESI) (Figure S3).

Finally, the 3 circular compartments are closed using PDMS lids to avoid evaporation (see Figure 5c), that were punched with Biopsy punches (Kai-Europe GmbH, Germany) to hold the pH electrodes and tubing for the pH adjustment and the gradual addition of simulated gastric fluids and enzymes. These lids are easy to remove or place back and thus enable easy sampling from the digestion chambers at any time using a standard micropipette.

### Temperature and pH sensing and control

Digital temperature sensors 2x2 mm (MAX30205 Human Body Temperature Sensor, Maxim IntegratedTM, San José, California, USA) were integrated at the bottom of the gastric and intestinal chambers (Figure 5c). The sensors have high accuracy (±0.1ºC) in the human body temperature range and have a small footprint. The temperature readings are used to modulate the power delivered to the heating elements, more precisely insulated Nichrome (NiCr) wires having high resistivity, via a proportional integral derivative (PID) control loop to maintain temperature constant at 37 <u>º</u>C.

For pH sensing, potentiometric Iridium oxide (IrOx) sensors were used. The electrodes were fabricated following the anodic deposition approach reported by Yamanaka et al.^34^, which yields hydrated IrOx films with high (super-Nernstian) pH sensitivity^35^. The metal oxide sensors have number of attractive characteristics for integration into microsystems, such as their simple construction and ease of miniaturization, high mechanical and chemical stability and low cross-sensitivity to various salts^36^. In our system the sensors were fabricated by electrochemical deposition onto a titanium wire substrate. IrOx sensors are also biocompatible and have even been shown in vivo applications^37^.

As reference electrode a simple single-junction Ag/AgCl reference electrodes encapsulated with 3M KCl solutions in micropipette tips were fabricated using an approach adapted from Barlag et al.^38^. In our system, we replaced the salt bridge with a Ca^2+^cross-linked alginate gel instead of agarose. A detailed protocol for the electrode fabrication is provided in ESI.

The electric potential difference between the pH electrodes is registered by an electronic circuit based on the ESP32 microcontroller. The real-time pH readings are used to control the pumping of 1M HCl or 1M NaOH from two inexpensive, custom-made programmable syringe pumps, in order to adjust the pH to the values set for each phase of digestion.

A Printed Circuit Board (PCB) was designed to interface the syringe pumps, temperature sensors, heating elements and mixer motors with an Arduino Uno microcontroller and other extra electronic components required for the automation. A conceptual schematic of the system can be found in ESI (Figure S4).

### *In-vitro* digestion

Detailed information on material references and suppliers can be found in ESI. Casein labelled with a pH-insensitive red fluorescent dye BODIPY TR-X (EnzChek™ Protease Substrate, 589/617 nm)^39^ and lipase substrate labelled with a green fluorescent dye BODIPY-C12 (EnzChek™ Lipase Substrate, 482/515 nm)^40^ were used to measure the digestion rate and study the enzyme kinetics during in-vitro digestions. These reporter molecules are quenched by proximity, but become fluorescent after exposure to digestive enzymes.

The simulated digestive fluids were prepared following the INFOGEST protocol (see ESI - Table S1 for detailed recipes) and stored at -20 °C before use. Enzyme activity of pepsin and pancreatin was determined according to the procedure described by Brodkorb et al.10. Samples of 100 μL in the static digestions and 20 μL in the semi-dynamic digestions were collected throughout the gastric and intestinal phases and enzymes were inactivated by adding Pefabloc® (5 mM final concentration in the digesta) to inhibit protease activity and Orlistat® (1 mM final concentration) to inhibit lipase activity. The digestion rate was quantified by measuring the fluorescence of the test molecules using a BioTeK® Synergy H1 microplate reader (Winnoski, VT, USA).

### INFOGEST static protocol

*In vitro* static digestions were carried out in a thermomixer (Eppendorf™, Hamburg, Germany) or inside a single compartment of the digestion-chip. Briefly, 1 mL sample (protease and lipase substrate) was mixed 1:1 with simulated salivary fluid (SSF) and incubated for 2 minutes at 37 °C. Following this, the simulated gastric fluid (SGF) and pepsin (final concentration 2.000 U/mL), were mixed 1:1 with the bolus, the pH adjusted to 3.0 (using 1 M HCl) and allowed to digest for 2 hours. Then, the simulated intestinal fluid (SIF) containing bile salts (10 mM) and pancreatin (final concentration of trypsin 100 U/mL) were added to the end-point of the gastric phase (chyme), the pH was increased to 7.0 (using 1 M NaOH) and the chyme was incubated for further 2 hours.

### Semi-dynamic protocol

The digestion-chip aims to faithfully emulate the *in vivo* conditions by incorporating some dynamic features, in the gastric phase, such as gradual acidification, addition of enzymes and simulated gastric fluid, and performing different gastric emptying. The semi-dynamic protocol followed here was adapted from Mulet-Cabero et al.^17^ and is represented in Figure 6. Glucose (100 mg·mL^-1^) was added to the enzyme substrate to increase the overall caloric content of the sample as the total incubation time in the gastric phase is calculated as a function of the caloric content of the sample, according to the spreadsheet provided by Mulet-Cabero et al., leading to 80 minutes of total gastric incubation time. Briefly, 100 mg·mL^-1^ glucose was added to 500 μL of substrate (protease and lipase substrates) and mixed 1:1 with SSF and then incubated for 2 minutes at 37 °C, in the gastric chamber. The gastric phase initiates with just 10% of the total amount of gastric secretions to be added owing to the fasting levels. Thus, 10% of simulated gastric fluids (100 μL of SGF + pepsin) was added to the oral phase in the gastric chamber and the remaining 90% (900 μL of SGF + pepsin) was gradually added using a New Era NE-2000 syringe pump (New Era Pump Systems, Inc., NY, USA) at a constant flow rate of 11.25 μL·min^-1^, achieving a 1:1 volume ratio with respect to the oral phase at the end of the gastric phase (80 minutes). The pH was gradually acidified from ≈7 to ≈2 at the end. This was achieved by a constant communication between the pumping of HCl (1M) using a custom-made syringe pump and the readings from the pH sensors. The gradual addition of HCl contributes to an increasing enzyme activity over the gastric phase.

During the incubation time in the gastric phase, sequential emptying into the intestinal chamber were performed. The number of gastric emptying is defined by the user, and in this case 4 were done, every 20 minutes. The gastric emptying was performed by actuating the peristaltic micro-pump, previously described. Chyme from the gastric chamber is mixed 1:1 with simulated intestinal fluids (SIF + bile salts + pancreatin). Here, 250 μL of chyme was mixed with 250 μL of simulated intestinal fluids after each emptying and incubated in the intestinal chamber for 2h. Following each emptying the pH was adjusted to 7.0 adding NaOH (1M).

## Acknowledgements

This work was funded by the SbDtoolBox - Nanotechnology-based tools and tests for Safer-by-Design nanomaterials, with the reference NORTE-01-0145-FEDER-000047, funded by Norte 2020, North-Regional Operational Programme under the PORTUGAL 2020 Partnership Agreement, through the European Regional Development Fund (ERDF) and by the European Union’s Horizon 2020 research and innovation programme through the project GASTRIC, under the Marie Sklodowska-Curie grant agreement no. 101003440.

## Author Contributions

**V. Calero:** investigation, methodology, data analysis, writing – original draft preparation, and writing – review and editing. **P. Rodrigues:** investigation, methodology, data analysis, writing – original draft preparation, and writing – review and editing. **T. Dias:** investigation, methodology, data analysis. **A. Ainla:** investigation, methodology, data analysis, writing – original draft preparation, and writing – review and editing. **A. Vilaça:** investigation, methodology, data analysis. **L. Pastrana:** writing – review and editing, funding acquisition, project administration. **M. Xavier:** conceptualisation, data analysis, supervision, writing – review and editing, funding acquisition, project administration. **C. Gonçalves:** conceptualisation, data analysis, supervision, writing – review and editing, funding acquisition, project administration.

## Data availability

The data generated during the current study is available from the corresponding author on reasonable request.

