## Supplemental Information for "A miniaturised semi-dynamic *in-vitro* model of human digestion"

### Electronic Supplementary Information (ESI): A miniaturised semi-dynamic *in-vitro* model of human digestion

#### Materials

Pepsin from porcine gastric mucosa, bovine bile, pancreatin from porcine pancreas were obtained from Sigma-Aldrich (St. Louis, MO, USA). EnzChek™ Protease Assay Kit and EnzChek™ Lipase Substrate were purchased from Thermo Fisher Scientific (Waltham, MA, USA). All reagents were used as received and according to the manufacturer's recommendations.

*Table S1. Concentrations (in mM) of electrolytes in the stock (1.25x concentrated) simulant digestion fluids used for digestion. The concentrations of $CaCl_2(H_2O)_2$ and bile salts indicated are that of the final digestion mixtures.*

| | Simulated salivary fluid (SSF) | Simulated Gastric Fluid (SGF) | Simulated Intestinal Fluid (SIF) |
|---|---|---|---|
| **KCl** | 15.1 | 6.9 | 6.8 |
| **$KH_2PO_4$** | 3.7 | 0.9 | 0.8 |
| **$NaHCO_3$** | - | - | - |
| **NaCl** | 13.6 | 72.2 | 123.4 |
| **$MgCl_2(H_2O)_6$** | 0.15 | 0.1 | 0.33 |
| **$(NH_4)_2CO_3$** | 0.06 | 0.5 | - |
| **$CaCl_2(H_2O)_2$** | 0.75 | 0.075 | 0.3 |
| **HCl** | 1.1 | 15.6 | 8.4 |
| **Pepsin** | - | 2,000 $U \cdot mL^{-1}$ | - |
| **Pancreatin** | - | - | 100 $U \cdot mL^{-1}$ |
| **Bile salts** | - | 10 | - |

#### Fabrication of pH electrodes

##### a. Reference electrode fabrication – Ag/AgCl electrode encapsulated in 3M KCl

Materials

- 0.1 M HCl solution
- Sodium alginate solution (0.5%)
- CaCl2 solution (3g/100mL)
- Ag wire (0.5 mm diameter)
- Pt wire reference (0.2 mm diameter)
- Cotton
- 200 µL pipette tip
- 3M KCl

Electrochemical deposition with 10mA/cm$^2$ for 1 min

- Place a small piece of cotton at the end of the pipette tip
- Fill the area with the cotton with 20-30 µL of the alginate solution
- Add ≈10 µL of CaCl2 solution from the top of the tip to the alginate solution
- Dip the bottom of the tip for at least 15 minutes in the CaCl2 solution

- Ag wire electrochemical deposition with AgCl
  - Place the Pt reference electrode and 1.5 cm length of the Ag wire inside the 0.1M HCl solution
  - Connect Ag to signal and Pt to ground to a DC power supply.
  - Apply a fixed current of 2.355 mA for 1 minute – the exposed area of the Ag wire must turn black (indicating deposition of AgCl)
  - Rinse the wires with DI water
- Fill the pipette tip with 3M KCl once the alginate gel is cross-linked – ensure there is no leakage from the bottom of the tip
- Place the Ag/AgCl wire inside the pipette tip immersing only the coated AgCl area in the 3M KCl solution
- Seal the top of the tip with parafilm

#### b. pH sensing electrode fabrication – IrOx deposition in Ti wire

##### Materials

- Solution for Iridium Oxide deposition (for 10 mL of volume)
  - Dissolve 15 mg of Iridium chloride hydrate ($IrCl_4.H_2O$) in 10 mL of DI water (magnetic stir for 30 min)
  - Add 100 µL of aqueous 30% H2O2 and stir 10 min
  - Add 50 mg of oxalic acid dehydrate ($(COOH)_2.2H_2O$) and stir 10 min
  - Adjust the pH of the solution slowly to 10.5 by adding small portions of anhydrous potassium carbonate (K2CO3) – resulting solution must be yellow
  - Cover the solution (to protect it from light) and leave it at room temperature for 2 days to stabilize. When the colour of the solution changes from yellow to light-violet it is ready to be used for deposition
  - The solution must be kept in the dark at 4ºC until is used – it preserves well for a couple of months
- 0.5 M $H_2SO_4$ solution
- Pt wire reference (0.2 mm diameter - Sigma Ep1330-1EA)
- Ti wire for IrOx deposition (0.25 mm diameter Sigma 460400-2.2G)
- 200 µL pipette tip and epoxy resin for encapsulation of bare Ti surface.
- 5% Nafion solution.

##### Electrochemical deposition with 1 mA/cm$^2$ for 20 min

- Fill the pipette tip with epoxy resin and insert the Ti wire before it cures. Leave a length of 10 mm exposed.
- Clean the electrodes by Cyclic Voltammetry – connect the Ti wire to the signal and the Pt reference to the ground. Apply a -0.23V to +1.1V at 100mV/s rate for 20 cycles in the $H_2SO_4$ solution (0.5 M).
- When the cleaning is completed rinse both wires with DI water and dry.
- With the same connections, immerse the wires in the Iridium Oxide deposition solution – Apply a constant current of 1 mA/cm$^2$ (specific for the area exposed – 10 mm length and 0.25 mm diameter of the Ti wire) for 20 minutes. After the deposition the exposed area of the Ti wire should turn dark blue.
- Rinse the wires with DI water and dry with N2.
- Dip coat the coated IrOx area in 5% Nafion solution and place in the oven at 65ºC for 20 minutes. Repeat the Nafion coating a second time.
- Store in a buffer solution similar to that for which it will be used for. Allow the solution to stabilize for 2 days.

#### c. Electrode storage

- In the stabilization solution for the IrOx electrodes.
- In 3M KCl for the encapsulated AgCl electrodes protected from light.

-

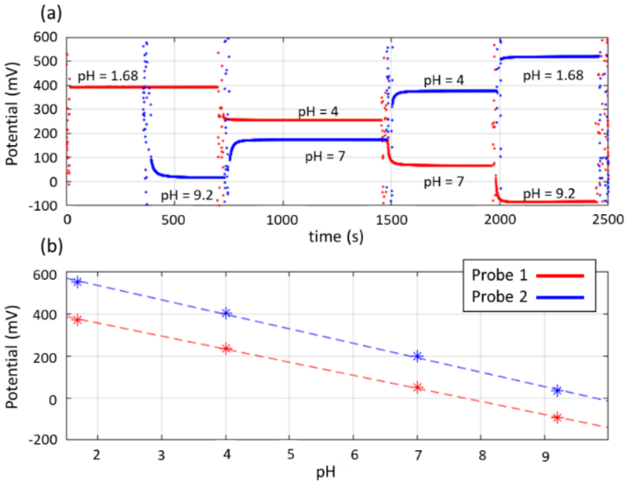


Figure S1. Example of the calibration procedure of two pH probes. (a) Response and stabilisation of the electric potential difference of two pH probes (blue and red) in standard pH calibration solutions. (b) Calibration curves showing the linear response of the pH probes.

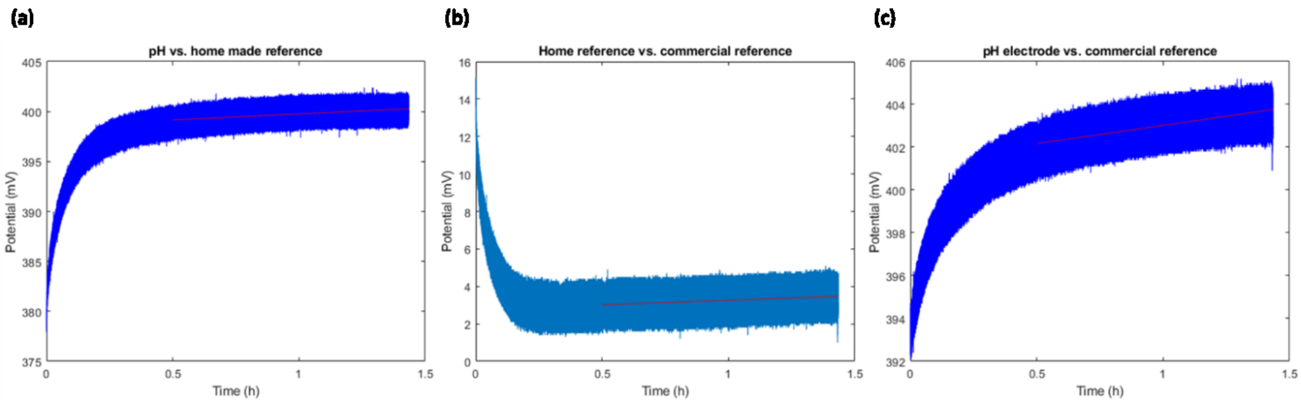


Figure S2. Electric potential stability tests of the pH electrodes in a pH=4 standard calibration solution. (b) Pair of pH electrodes fabricated in-house. (a) Fabricated reference electrode vs commercial probe reference. (c) IrOx sensing electrode vs commercial probe reference.

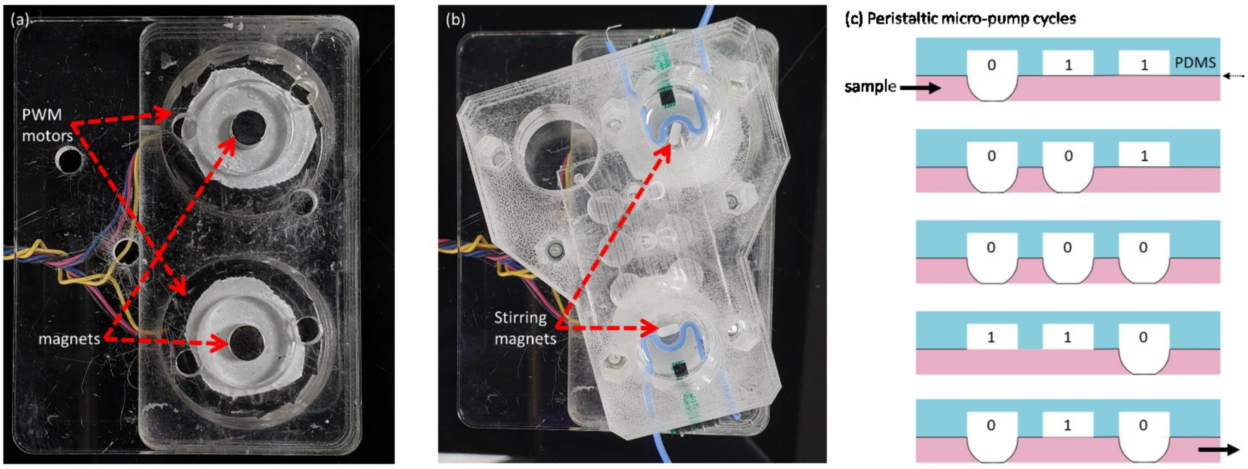


Figure S3. Description of the stirring system for efficient constant mixing inside the digestion chambers. (a) PWM motors that allow to control and reduce the mixing speed to avoid enzyme degradation. (b) Small 5x2 mm stirring bars placed inside the digestion chambers, which are driven by the motors placed underneath. (c) Depiction of the peristaltic micro-pump cycles.

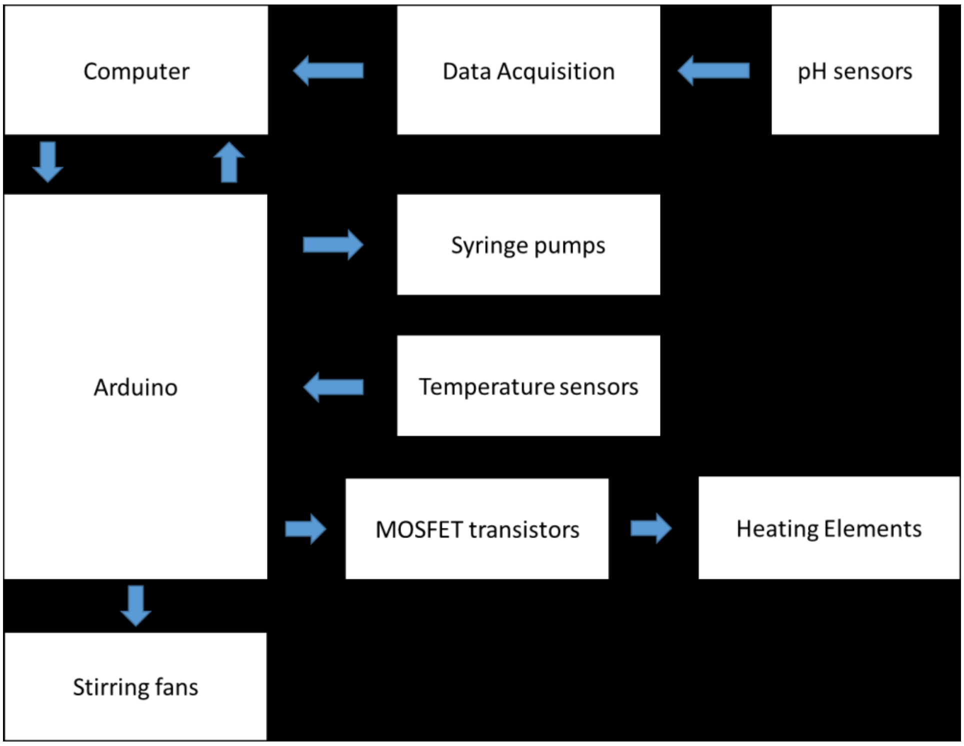


Figure S4. Description of the circuit for automated operation of the digestion system. The heating elements and stepper motors (SparkFun Electronics) of the syringe pumps were powered using a 12V power supply. The pH data acquisition electronic board was based on a ESP32 microcontroller.
